# Negative epistasis limits current codon optimization approaches

**DOI:** 10.1101/2025.06.03.657573

**Authors:** Marco Gees, Zrinka Raguz Nakic, Maria Anisimova, Victor Garcia, Christin Peters

## Abstract

Demand for high-yield protein production in biotechnological applications is driving efforts to maximize heterologous protein expression in scalable microorganisms such as *E. coli*. While codon optimization techniques employed by contemporary sequence providers promise high-expression products, expression levels are often unsatisfactory. Whether the causes for this performance unreliability are due to fundamental constraints on the predictability of protein yields, or whether they stem from differences in theoretical approaches, is unknown. Here, we performed a comparative analysis to address this question. We assessed the performance of twelve different optimization approaches at enhancing expression of a sequence encoding a cinnamyl alcohol dehydrogenase. Six approaches stemmed from commercial providers and six from freely available sources. Through analysis of their elongation time profiles and multidimensional scaling we assessed which algorithms follow a unique optimization approach. We found that codon-optimized sequences are, on average, capable to raise protein expression levels with respect to the nonoptimised source sequence. However, variation in the protein expression levels was large. Simple, non-proprietary optimization techniques were capable of achieving protein expression levels that fall within the top expression range amongst candidate sequences. Lastly, we found that negative epistasis influences a sequence’s protein expression level. Since therefore the protein expression landscape arising from synonymous sequence space must exhibit a non-negligible degree of ruggedness, standard approaches will be limited in their capacity to predict protein expression levels of sequences.

## Introduction

The increasing demand for high-yield proteins in therapeutic [1], industrial, and research settings drives the need to harness their expression at significant scales [2]. A prominent strategy in this domain is heterologous expression, where a protein native to a particular ‘origin’ organism is produced in a ‘target’ organism that can more easily be scaled for mass production [3, 4].

Heterologous expression approaches face significant challenges. Genomic sequences from one organism have evolved over large time spans, and have been tailored by evolution to match the unique genomic environment of their host [5, 6]. Such sequences are replete with modifications that reflect the past environmental pressures faced by the source organism. Given the great energetic costs imposed on the organism by translation [7], sequences are expected to reflect, in particular, selection for cost efficient translation [5, 8, 9]. Translocating these sequences into a different host organism introduces them to a foreign genomic landscape, characterized by new and different translational features. Thus, selectively beneficial changes acquired in the source context might turn maladaptive in the new target environment, jeopardizing yield [3, 4, 10, 11].

A common approach to mitigate these risks is *codon optimization*, a method that alters the source sequence composition via synonymous substitutions, thereby preserving the order of amino acids that it encodes [12–14]. Different codon optimization approaches and algorithms are currently made available –especially for *Escherichia coli* – by a variety of commercial providers of transgenic DNA products as well as open-source software [15]. However, the reliability if these algorithms is generally poor [16] and the source codes of many of the algorithms in use remain undisclosed, complicating a comprehensive assessment of current-level capacities to predict protein expression yield for a sequence.

In this study, we investigated the performance of six current codon optimization techniques by commercial providers as well as six non-proprietary optimization algorithms for the *A. thaliana* gene CAD5 (AtCAD5), which we heterologously expressed in two strains of *E. coli*, BL21(DE3) and K12(DE3). Our study is divided in three parts. First, a characterization of the sequences themselves and their inter-relationships according to the optimization approach employed. Second, the description of the protein expression features of these sequences. And third, we combine the information of these two assessments to test for the presence of epistasic effects in protein expression.

We conclude that the observed variability in the expression levels attained may partly be explained by the inherent ruggedness present in the protein expression profile of the synonymous sequence of the AtCAD5 sequence. This result implies that there exist fundamental limitations on the predictability of protein expression levels of sequences by current approaches.

## Results

### The landscape of codon-optimized sequences

To assess the performance of currently used codon-optimization techniques, we heterologously expressed 22 distinct sequences of the *AtCAD5* gene, originally described in *Arabidopsis thaliana*, in two common laboratory strains of *E. coli*, BL21(DE3) and K12(DE3). AtCAD5 encodes a catalytically active cinnamyl alcohol dehydrogenase, a protein class which is expected to express well in *E. coli*. In plants, it is involved in and acts upstream of the lignin biosynthetic process (see Table II for references and details), which has received increasing attention for its potential to sequester carbon and thus mitigate the effects of global warming [17, 18]. Two splice variants are known, of which we chose AT4G34230.1, the longer variant, whose coding sequence contains 357 codons.

**TABLE I.**
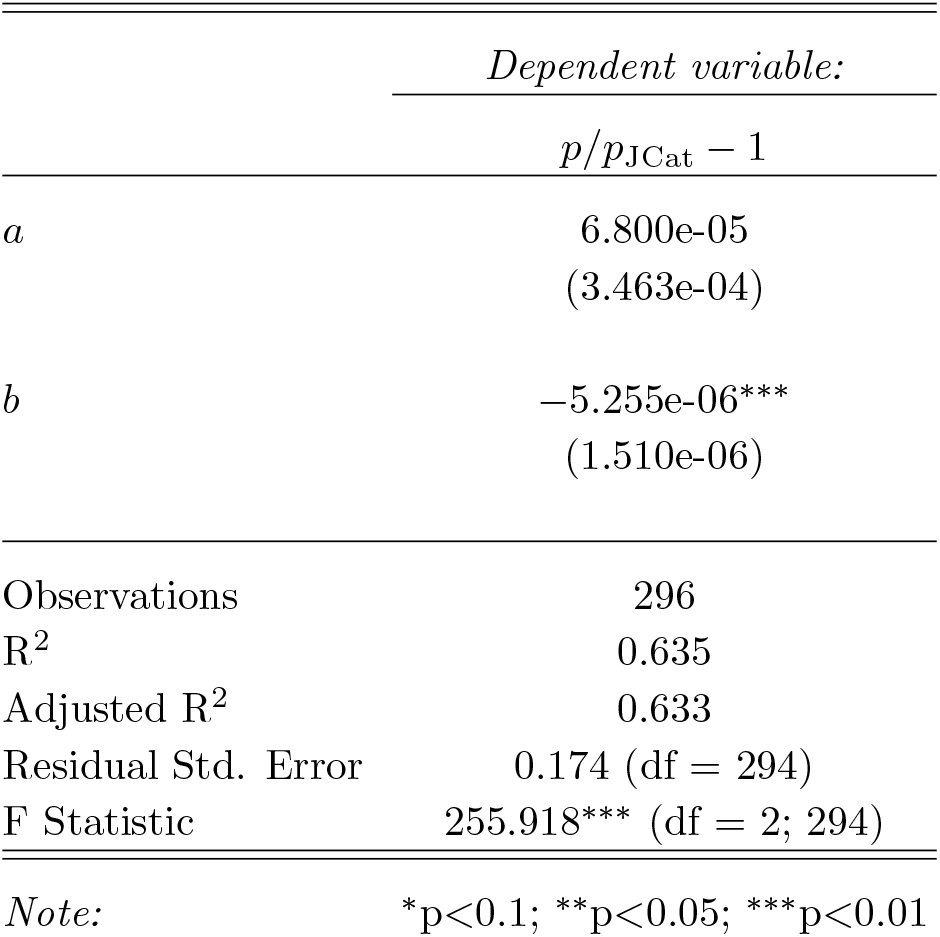

| *Dependent variable:* |  |
| --- | --- |
| | $p/p_{\text{JCat}} - 1$ |
| $a$ | 6.800e-05<br>(3.463e-04) |
| $b$ | -5.255e-06***<br>(1.510e-06) |
| Observations | 296 |
| R<sup>2</sup> | 0.635 |
| Adjusted R<sup>2</sup> | 0.633 |
| Residual Std. Error | 0.174 (df = 294) |
| F Statistic | 255.918*** (df = 2; 294) |

*Note:* \*p<0.1; \*\*p<0.05; \*\*\*p<0.01

**TABLE II:**
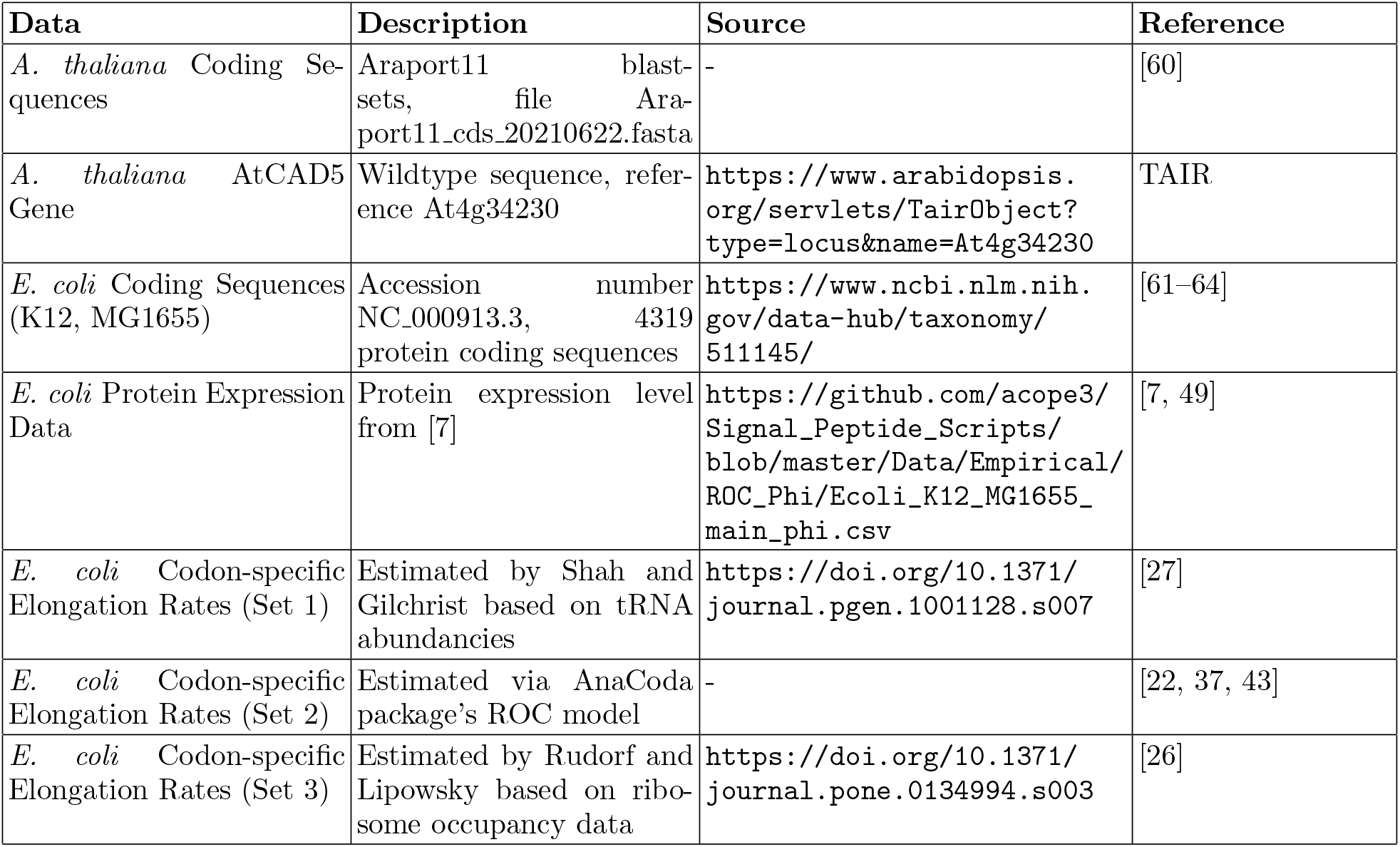
Summary of data sources used.

These 22 synonomous sequences were optimized by either proprietary software, open-source algorithms or by algorithms developed at the Institute of Computational Life Sciences (ICLS) in Wädenswil. In the proprietary subset, we used optimization algorithms from *Twist Bioscience*, the *GENEius* algorithm by *Eurofins Scientific SE, Genscript Biotech, NovoPro Bioscience, GeneWiz* from *Azenta Life Sciences* and *Integrated DNA Technologies* (IDT) (see Materials and Methods for overview and Supporting Information for detailed sequence description). Three sequences were produced by opensource algorithms: One sequence was optimized using the package *GeneGA* [19] within the Bioconductor package [20], one with JCat (Java Codon Adaptation Tool) [21], and one with the use of the AnaCoda package [22, 23].

Another three sequences were optimized by algorithms developed at the Institute of Computational Life Sciences at ZHAW. The rationale behind the development of our own optimization strategy was based on the notion that indiscriminately pruning sequences from translationally inefficient codons can jeopardize the stability of the translational and folding processes [24, 25]. Where these codons are preserved by purifying selection, they are likely supporting a key function for the organism, and should therefore be retained. To implement this idea, three approaches were developed: A *mirrored* approach, where codon efficiencies in the host sequence are matched to the codon efficiencies of the source sequence. In a second approach, termed *mirrored and restricted* the sequence is additionally pruned of a predefined set of restriction sites. In a third approachwe replaced the most inefficient codons from the *mirrored and restricted* sequences with the second-most inefficient codons, giving rise to the *mirrored and capped* sequence candidate. A summary of the set of sequences used and the references to their design algorithms is given in Table III.

**TABLE III:**
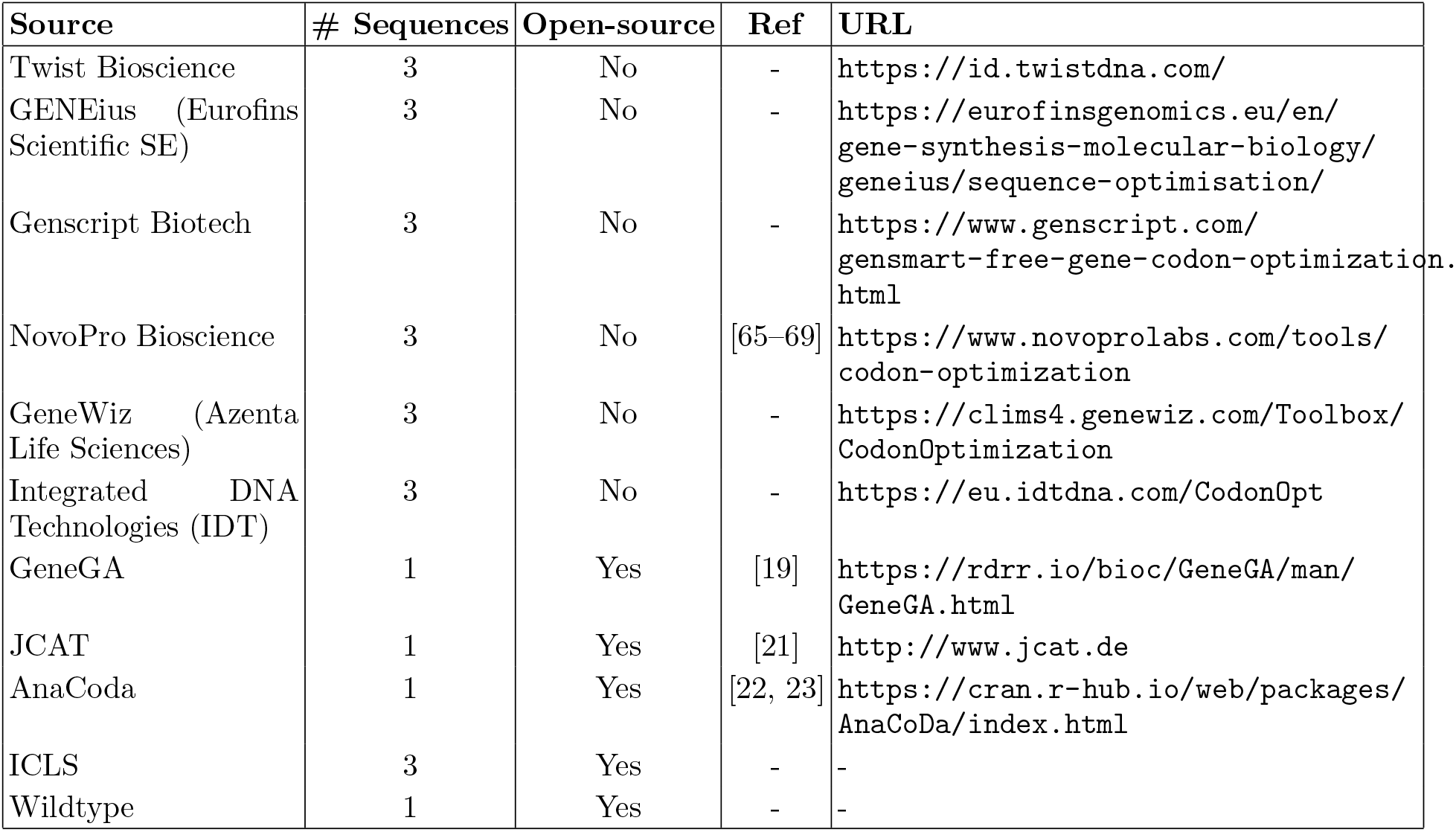
Summary of the heterologous expression of the AtCAD5 gene sequences in E. coli. The table lists the sources or algorithms used for codon optimization, the number of sequences optimized by each source, whether the generating algorithm is open-source, references where the algorithm is describe in detail (where available), and the urls where the optimization algorithms were used or may be downloaded.

All sequences were designed with a C-terminal tag that consists of 30 codons that encode for a linker and GFP11 (See *Supplementary Information*). The tag is used for a Split-GFP assay where it complements to an intact GFP upon addition of the detector fragment GFP1-10, resulting in fluorescence that is proportional to the amount of the expressed protein, thereby allowing quantification of the expressed protein.

The optimized sequences display variation in the number of synonymous changes made. Relative to the wildtype, the sequences from Twist Bioscience show the largest number of point substitutions, with about 35% of the original sequence altered. These substitutions translate to about 92-94% of codons changed. The rest of the optimization algorithms result in smaller modifications. About 22-26% of the sequence is altered by point substitutions by the rest of the optimizers, corresponding to 60-67% of codons changed. Thus, sequences optimized by Twist Bioscience are unique in the amount of modification they undergo.

Prior to analyzing the expression features of these sequences, we first investigated how they differ in their efficiency profile, and second, how they are distributed in sequence space.

Figure 1 shows the local elongation time profiles of all optimized sequences for three different sets of codon-specific elongation time estimates, all inferred via distinct theoretical approaches. The first set of elongation time estimates was inferred from genome and protein expression rate data with the AnaCoda package [22, 23] (Fig. 1A). A second set of estimates was inferred from ribosome occupancy data by Rudorf et al. [26] (Fig. 1B). The third set was inferred from tRNA abundance data by Shah and Gilchrist [27] (Fig. 1C), see *Materials and Methods* for detailed descriptions). For all three sets of elongation time estimates, the vast majority of approaches for codon optimization decrease the average time of translation relative to the original wildtype sequence, or equivalently, raise the average translational rate per codon.

**FIG. 1:**
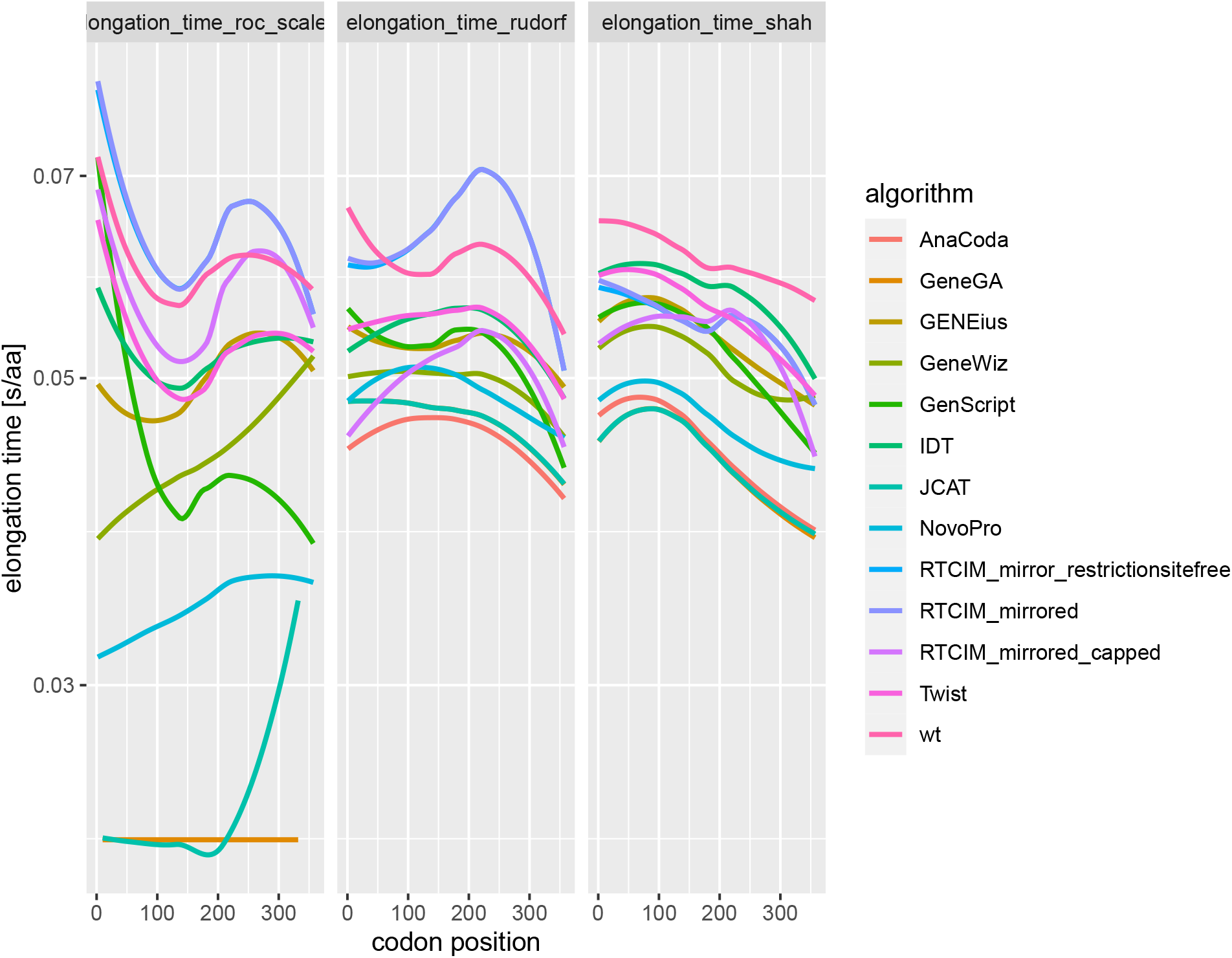
Translation efficiency profiles by optimization algorithm. Locally estimated scatterplot smoothing (LOESS) curves of the codon-specific relative elongation times (y-axis) for all three sets of elongation time estimates (panels A to C) by codon position (x-axis). A: LOESS Curves for the AnaCoda-derived relative translation inefficiencies - a proxy for elongation times-given relative to the optimal codon within a family. B: LOESS-curves of efficiency profiles based on the Rudorf et alḋata set [26]. C: LOESS curves for the efficiency profiles based on the Shah and Gilchrist codon-specific elongation time estimates [27]. For algorithms with multiple optimized sequences, the elongation time profiles were aggregated prior to running the LOESS regression due to low within-algorithm sequence variation.

The efficiency profiles of the optimized sequences reflect the different strategies used by the algorithms. All open-source based strategies follow an all-optimal codons approach, but with differences in the identity of the optimal codons. The exception are the ICLS-based sequences, which duefully replicate local elongation time patterns of the wildtype. In contrast to the open-source algorithms, proprietary algorithms do not solely follow an all-optimal codons strategy, as evidenced by their deviation from the AnaCoda and JCat sequences, which in fact do follow such a strategy. Instead, maximum efficiency codons are frequently traded off for codons that optimize other, undisclosed features - most likely features linked to the probability of obtaining viable protein. The wildtype sequence has among the highest average elongation times of all sequences –in line with expectation– together with some instances of the ICLS-based sequences.

We proceeded to utilize these efficiency profiles to characterize the inter-sequence relationships among the optimized sequences via Multidimensional Scaling (MDS, see *Materials and Methods*).

Figure 2 shows the MDS of all the optimized sequences computed with the elongation time (for AnaCoda-based data, Figure 2A) and elongation rate (for Shah and Rudorf data, Figures 2B and C) profiles of these sequences as derived from all three codon-specific elongation time data sets. Additionally, panel D shows the MDS derived from the Hamming distance as a similarity metric. The different codon-specific elongation time data sets give rise to qualitatively similar graphs. Sequences produced by the same algorithm tend to cluster together for all three data sets. The ICLS-based sequences, the Twist sequences and the wildtype produce isolated clusters, indicating idiosyncratic underlying design principles. The non-proprietary sequences are found at the periphery of the graphs, nearby a larger cluster of the rest of the sequences in which boundaries between algorithms are more fluid.

**FIG. 2:**
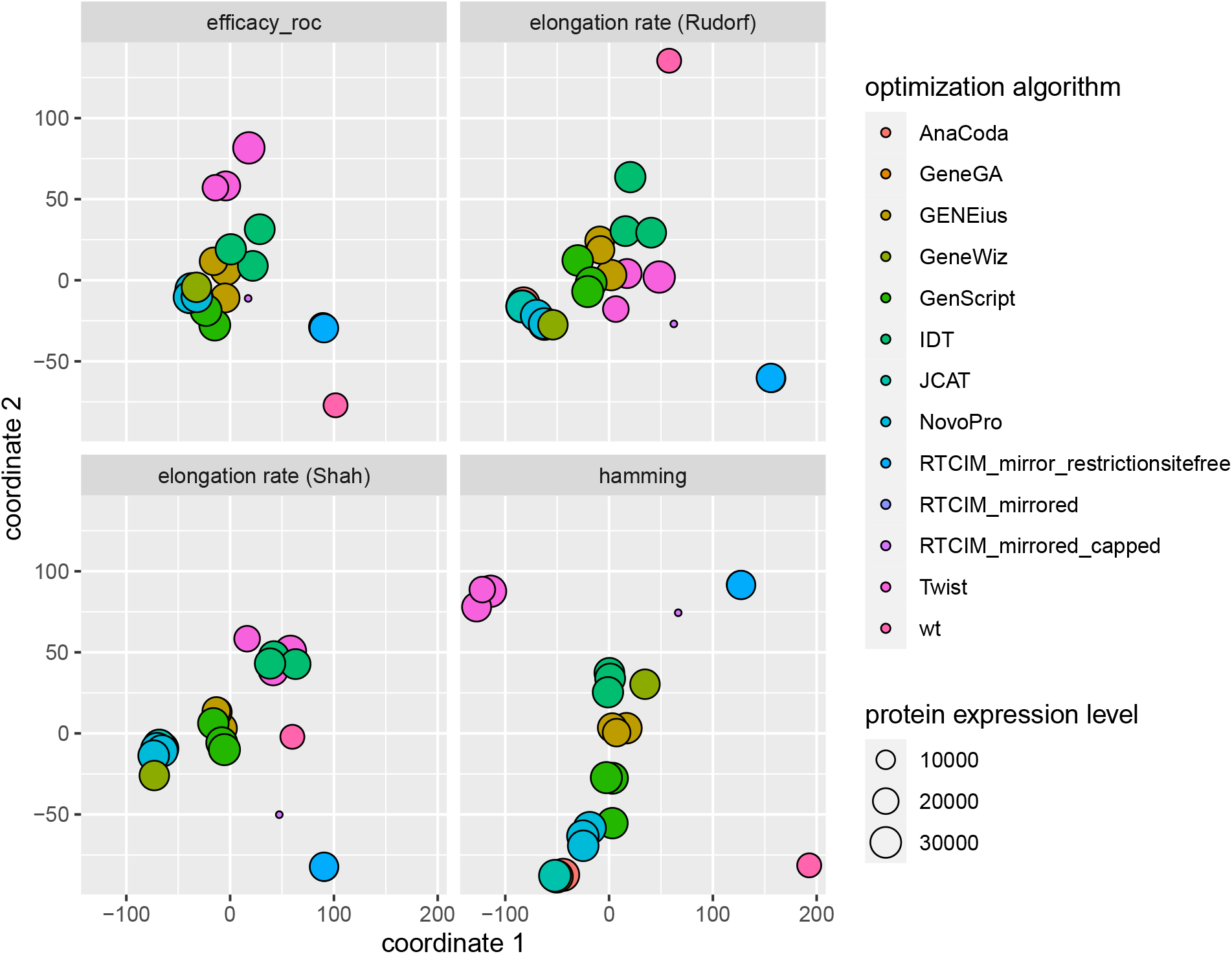
Multidimensional scaling for four different distance metrics between sequences. For A, the stress function after MDS optimization based on sequence distances of AnaCoda-derived elongation times was 16.8. For B and C, sequence distances were based on elongation rate profiles (Shah and Rudorf data, respectively), and the stress values were 34.8 and 23.4, respectively. D Hamming distances between sequences were used as similarity metric (stress: 35.8).

The distinctions discernible among algorithms are much more pronounced when Hamming distances are taken as the similarity metric. In Figure 2D, most sequences are spread out along a line stretching from position (0,−100) to about (0,40). Single sequence clusters by the same algorithm remain discernible, but the clusters increasingly overlap for sequences close to position (0,−100). This suggests that the sequences’ positions relative to the wildtype in sequence space may be approximated as being spread out along a particular axis. Across all MDS outputs, no clear trend for increasing protein expression is identifiable along any direction.

These results suggest that the commercial providers do deploy optimization algorithms that give rise to distinct sequence types, but that these differences diminish in importance if elongation time or rate similarity metrics are adopted.

### Protein expression levels of codon-optimized *AtCAD5* sequences

Next, we assessed by how much the optimized sequences were able to raise protein expression levels relative to wildtype. To this end, all optimized sequence variants were expressed in BL21(DE3), a standard *E. coli* strain for biotechnological protein expression. To investigate the transferability of the findings to other strains, a subset of sequences was also expressed in K12(DE3), another standard research strain (from here on we term the strains BL21 and K12, respectively). To assess expression levels, a Split-GFP assay was performed, and relative protein amounts were determined by fluorescence signal [28]. For each optimized sequence, a total of eight replicates (two quadruplicates) were generated to measure fluorescence. Additionally, activity was determined in enzymatic assays using cinnamyl aldehyde as substrate.

The optimization methods reliably generate high-expression products (see Figure 3). For the BL21 strain, 95.5% of the optimized sequence replicates exceed the mean wildtype expression level. Only the *mirrored and capped* -strategy designed by the ICLS attains overall inferior expression levels than the wildtype. In the K12 strain, 91.3% of the replicates exceed the mean of the K12-wildtype. Here, it is one of the *Twist* sequences that yields lower average expression values than the wildtype. For the BL21(DE3) strain, the average fold increase over all optimized sequences was 1.61, whereas for K12 it is 1.96.

**FIG. 3:**
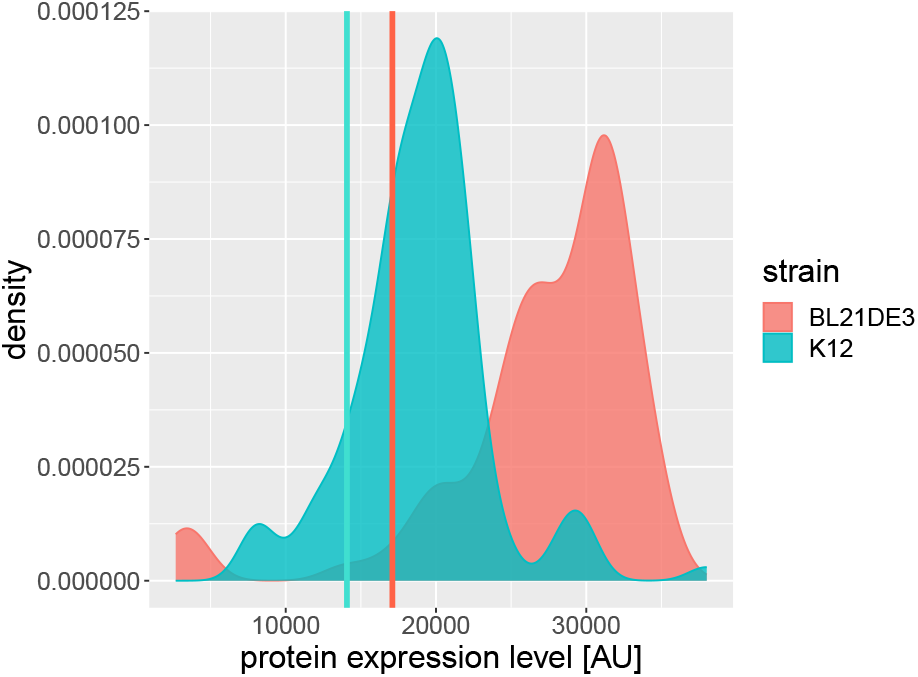
Empirical density distribution of the protein expression levels of all replicates of the 22 optimized sequences of BL21 and K12 strains. The red and green vertical lines represent the mean wildtype expression levels taken over all eight replicates for BL21 and K12 strains, respectively.

Figure 4A shows that the maximum mean fold increase of 1.95 for the BL21 is attained by the optimized sequence *NovoPro [1]*, designed by the Novo-Pro Bioscience. For K12, the maximum expression is attained by the *JCat* sequence, with 1.85 mean fold increase. The *mirrored and capped* strategy for the BL21 strain displays exceptionally low protein expression levels.

**FIG. 4:**
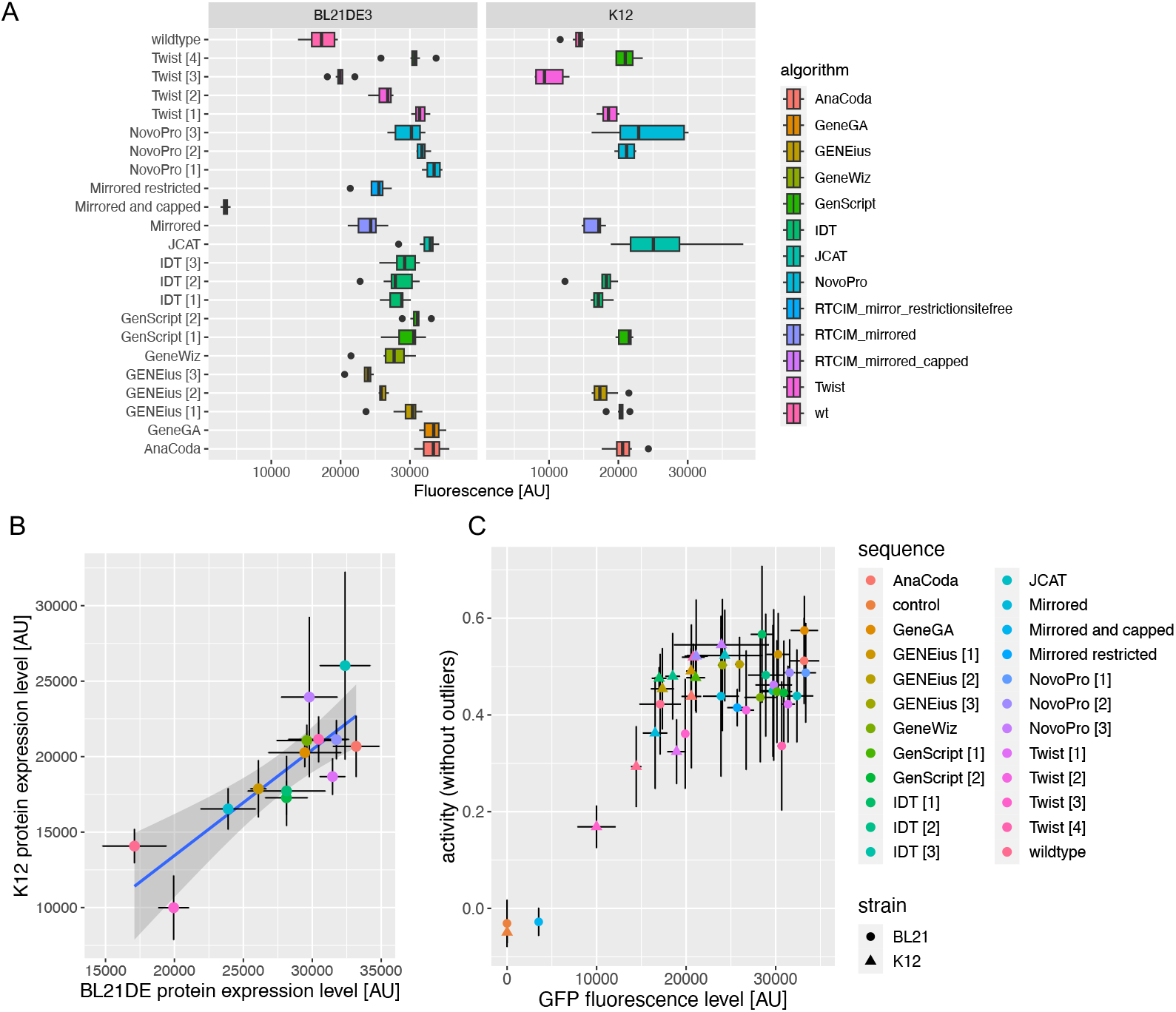
Protein expression characteristics of codon optimized sequences of the *AtCAD5* gene for BL21 and K12. (A) Protein expression levels (in arbitrary units (AU) of fluorescence levels) for all 22 optimized sequences expressed in the BL21, and for 13 sequences expressed in K12. (B) Protein expression levels of BL21 versus K12. Estimate uncertainties are given with two standard deviations (sd) –one sd for each direction pointing away from the mean– of eight replicate measurements taken per sequence and strain. Standard linear regression is shown as a blue line, while the grey area represents the 95% confidence range of possible regression line slopes under a Student t-distribution. (C) Protein expression levels versus enzymatic activity for expression products from both BL21 and K12. Uncertainty ranges are computed as in B).

Figure 4A reveals that for the BL21 strain, the intra-algorithm variation –that is, the variation in yield among sequences generated by the same algorithm– is generally small compared to the overall range of protein expression levels, as seen in the cases of NovoPro, IDT and Genscript, with the exception of the Twist-based sequences. For BL21, intra-algorithm variation is also larger in magnitude to intra-sequence variation –the variation in yield of eight replicates of the same sequence. Notably, all open-source algorithms reliably expressed very high protein levels, all close to the 33’000 AU from Novo-Pro [1], the most highly expressed sequence on average. Figure 4A also reveals significant stochasticity among the eight replicates assessed per sequence, especially for the K12 strain, mostly arising from inter-quadruplicate variation.

To test for the presence of strain-specific effects on expression, we computed the correlation of BL21 and K12 expression levels, finding a correlation of 0.83. Linear regression between expression levels across the two strains yields an estimated slope of 0.7, where the null that the slope equals zero is rejected with p-value < 0.001. This indicates that qualitatively as well as quantitatively, the expression characteristics of the sequences are very similar among the two strains.

The readouts from Split-GFP must be interpreted with care regarding functionality. Especially proteins at high expression levels are prone to improper folding and formation of inclusion bodies. To assess whether the expressed proteins are functional, we determined the enzymatic activity of the proteins generated by all sequences for BL21 and K12. To quantify the active enzyme, depletion of NADPH was monitored at A_340nm_ (absorbtion at 340 nm) upon consumption of the substrate cinnamal aldehyde. Figure 4C) shows that protein activity and expression levels are highly correlated across both strains BL21 and K12, with 0.8 overall correlation. Linear regression between expression and activity levels confirms the presence of a significant association with p-value < 0.001. However, closer visual inspection suggests the presence of an activity plateau after about 20’000 AU. This indicates that above a specific expression level, protein folding might become impaired, resulting in a partially non-functional protein pool.

In conclusion, we find that proprietary as well as open-source algorithms are able to consistently raise protein expression levels relative to the wildtype, with the exception of the ICLS-based *mirrored and capped* sequence in the BL21 strain and the *Twist [3]* sequence in the K12 strain. We find that the Split-GFP assay is well-suited to capture the protein expression landscape of the optimized sequences examined, with some limitations regarding assessment of protein functionality, and that this expression landscape retains its basic properties when expressed in a different *E. coli* strain. Our results however are also in line with previous observations that too high expression rates eventually result in non-functional protein presumably due to impaired protein folding, as shown in the enzymatic assays.

### Evidence for negative epistatic effects among synonymous codons on protein expression levels

Next, we addressed the question of whether the contributions from individual loci to overall protein expression combine in an additive, non-interactive manner, or whether alternatively, codons contribute non-additively, that is, whether they interact epistatically. To this end, we combined relational information on sequences with information on their measured expression. The methods for testing epistasis in these combined data is taken from theory devised to study epistasis in the context of population genetics.

For the purposes of protein expression maximization, one relevant insight from epistasis research is that the presence of epistatic effects in protein expression impedes the prediction of the protein expression level of a sequence – whereas if epistasis is low or absent, only relatively small numbers of sequences have to be measured to provide accurate predictions for unobserved sequences [29]. This is due to epistasis giving rise to a rugged sequence-to-expression landscape, while the lack of epistasis gives rise to a gradually rising landscape [29].

Several methods have been proposed to assess the presence of epistasis in population genetics contexts [29, 30]. One method, in particular, can be adapted to the data set of this study to address the question of whether codons in a sequence interact to affect the resulting protein expression level. The method is based on first determining a reference sequence, which is set to have zero log-fitness. This sequence is assumed to occupy a state of high fitness relative to its genomic surroundings. Log-fitness is then measured for sequences with deleterious mutations via regression,

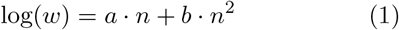

where *w* is the organism fitness, *n* is the Hamming distance to the reference strain and *a* and *b* are regression coefficients [30, 31]. A negative *b* coefficient is indicative of negative epistasis, while a positive value of *b* is interpreted as the presence of positive epistasis. Similar studies have been carried for relative short genomes in terms of log-fitness [32].

Since selection operates on the phenotype of an organism, the fitness of an organism is typically thought to receive contributions from a variety of traits of the organism. If such trait –such as protein expression level– is closely linked to fitness, it may be interpreted as an intermediate feature from which fitness is derived –in particular if all other traits are held constant in an experimental setting. Therefore, the methods and logic utilized to investigate epistatic effects on fitness are also applicable to the trait itself.

Thus, to test for epistatic interactions in the generation of the trait of protein expression, we adapted this method to the specifics of our data set. We applied the regression

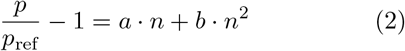

against the data. Here, *p* is the protein expression level, *p*_ref_ is the protein expression level of the reference sequence, and again, *n* is the number of deleterious mutations separating a reference sequence from any other sequence. The coefficients *a* and *b* are interpreted as above [30, 31].

To test this hypothesis, we first identified the best-performing sequence in both strains. The JCat sequence stands out as the one sequence which produces very high expression levels in both BL21(DE3) and K12 (see Figure 4B). As the JCat sequence is best performing sequence, the mutations that separate other sequences from it are assumed to exert a deleterious effect upon protein expression.

We utilized the Hamming distances computed for the multidimensional scaling computations between JCat and all other sequences to fit the regression formula (2) to the data. Figure 5 shows that the regression curve adopts a convex shape in both strains, BL21 an K12.

**FIG. 5:**
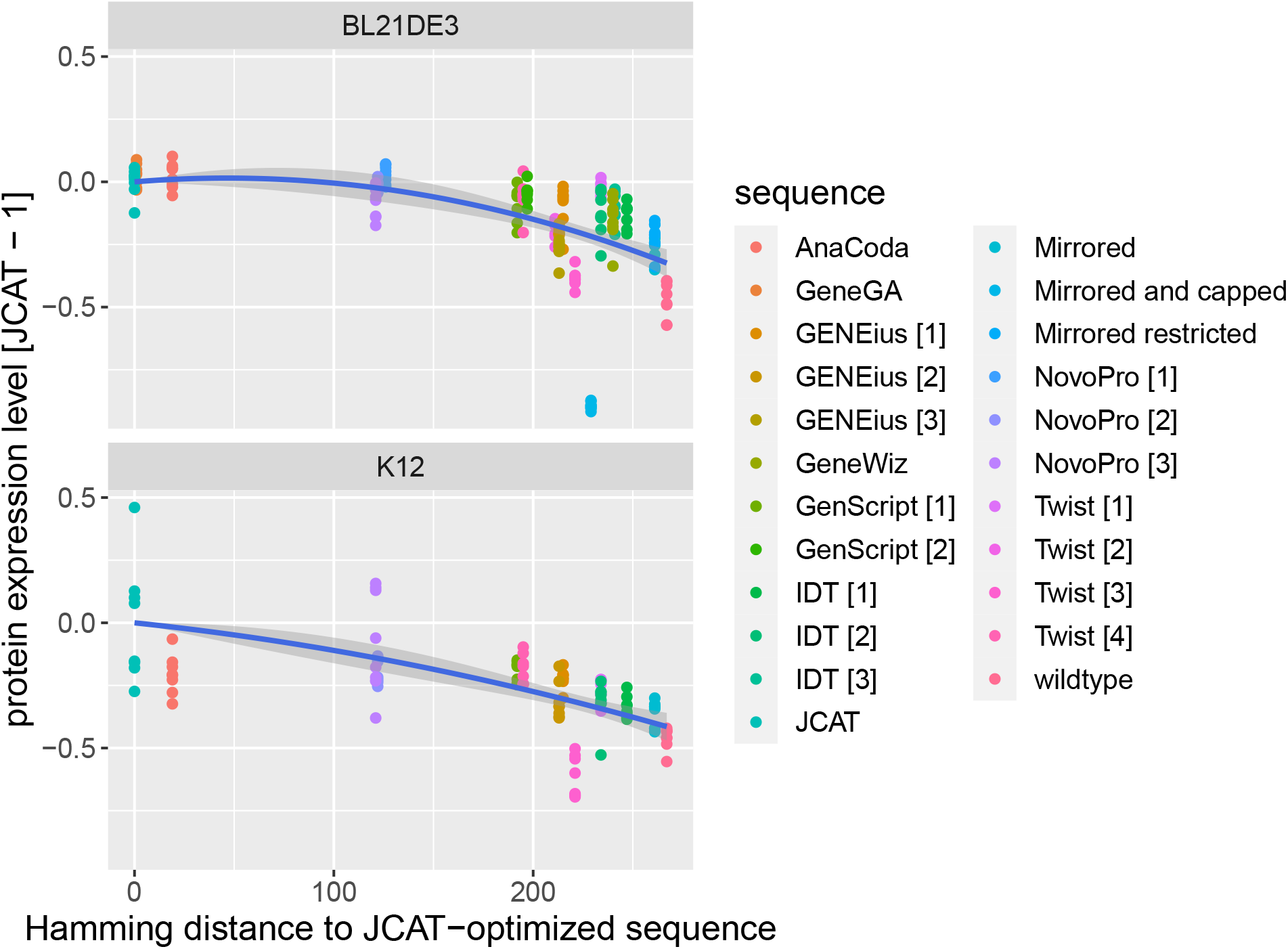
Negative epistatic effects among synonymous mutations combine to affect protein expression in BL21. Upper panel: Hamming distance from reference strain –the JCat optimized sequence– versus protein expression levels in units of the mean protein expression of the JCat sequence, minus one. Data points are shown for all measured replicates per sequence. No outliers were omitted. The fit of model (2) is shown as a royal blue line. The grey area around the regression line indicates the 95% uncertainty in the estimates of the two evaluated coefficients. Lower panel: Analogous to upper panel, but for the data measured in the K12 strain.

Fitting (2) on the combined data from BL21 and K12 yields a very strong statistical signal for a negative *b* value (see Table I, generated with the *stargazer* -package [33] in R [34]). This is indicative of negative epistatic interactions governing the generation of protein expression levels.

Running the same regression model, (2), on the strain-specific data sets individually yields more varied results. For BL21, the results of the aggregated data set are largely replicated. The statistical signal for a negative *b*-coefficient is strong, estimated at *b* = −7 · 10^−6^ with *p <* 0.001 for the null hypothesis H_0_ : *b* = 0. The adjusted R-squared for the regression is 0.54 at a number of observations of *n* = 184 (here, each replicate accounts for a data point). While the estimate for the *b* coefficient in the K12 data is negative (*b* = −2.7 · 10^−6^), it fails to attain statistical significance, most likely due to a limited number of observations (*n* = 112 versus *n* = 296 in the aggregated data set).

Summarizing, our data is consistent with the notion of negative epistatic effects governing the protein expression levels of the *CAD5* sequence. While individual analysis of the K12 data set fails to attain statistical significance for negative epistasis, all data sets that exceed K12’s number of observations do, raising confidence that the effect is present.

## Discussion

To express heterologous proteins at high yields, many biotechnological approaches today rely on a handful of microbial host organisms such as *E. coli*. However, optimization of expression for high yields is still a challenge. To address this issue, it is currently standard procedure to rely upon codon optimization, the synonymous modification of the original sequence to match the genomic and environmental features of the host organism.

Codon optimization is typically offered, free of charge, by most commercial sequence providers. The providers commonly claim that their optimization will –with high probability– result in functional protein at a high yield. However, comparative studies to examine these claims are scarce [11, 16].

The expectation that a model may be able to predict which synonymous sequences will yield high expression levels with accuracy is hard to justify on the grounds of the current body of theory on the mechanisms of translation. How codon usage bias in particular influences expression is determined by how synonymous changes affect translation, which remains subject to intense research efforts.

In fact, a plethora of factors related to synonymous sequence structure have been found to exert significant influences on protein expression [12]. Examples include the presence of codon ramps at the 5’-end of the sequence, mRNA secondary structure, tRNA abundances, microorganismal growth rates, GC-content, codon usage bias and so forth [11, 12]. These developments cast further doubts on the promise of accurate protein yield prediction from sequence information alone.

The predictive capability of any model aiming to derive protein expression levels from sequence structure relies on its capacity to capture the inherent features of a part of the genotype-phenotype map for that sequence [29]. The features of such a genotype-phenotype map are open to empirical study. These studies allow to ascertain whether the map displays features –for instance a pronounced inherent ruggedness– that preclude accurate prediction by means of standard approaches or not.

This study contributes to addressing this issue on three fronts. First, we validate that a main property of currently employed optimization algorithms is the reduction of total elongation time, or conversely, the increase of translational speed. We further represent our set of optimized sequences in via multidimensional scaling (MDS), and find that sequences of the same algorithm tend to cluster, confirming their common origin and high similarity. When using Hamming distances, MDS yields more pronounced algorithm-oriented clustering, indicating the presence of unique design principles. Second, we use a recently developed protein expression quantification technology, Split-GFP, for the purposes of genotype-phenotype map characterization on a protein that is not GFP itself. The technology yields measurements of high validity and reliability, as confirmed by further tests on the association of protein expression exhibited by two different *E. coli* strains. Third, we find that the synonymous genotype-to-phenotype landscape for protein expression is influenced by negative epistasis, and thus is very likely exhibiting ruggedness. This implies that fundamental limitations are imposed on the accuracy of protein expression prediction via naïve approaches by the nature of intra-sequence codon interactions. Thus, our results suggest that the study of the origin of epistatic effects on protein expression may help improve the predictive accuracy of protein expression models.

Our results come with a series of caveats. For instance, we have explored the expression landscape of a fraction of the possible synonymous sequences of a single heterologously expressed gene, AtCAD5. Whether the specific biochemical features of AtCAD5 or its protein product preclude the generalization of our results to other is hard to predict. However, most larger expression landscape studies of synonymous sequences focus on green fluorescent protein (GFP) only [35, 36]. Our study contributes to the investigation of the robustness of the GFP-derived conclusions by the exploration of another protein that is not GFP.

We also lack a firm explanation for the failure of a subset of sequences to attain similar expression levels as the rest of the sequences, notably the *mirrored and capped* sequence in BL21(DE3), and the *Twist [3]* sequence for K12. One possible reason is the formation of dense aggregates of inclusion bodies upon high expression, which do not only affect protein activity, but also mask the GFP tag. The formation of fluorescent GFP entities would be hindered in such cases, rendering the Split-GFP assay incapable to detect such aggregates. Furthermore, such unexpected declines of protein expression below the original sequence’s level, are consistent with a rugged synonymous genotype-phenotype map. Ruggedness implies the presence of “expression valleys”, that is, sequences or localized sequence clusters where expression suddenly falls off. The presence and location of such valleys is hard to pinpoint by means of codon usage bias theory alone.

Furthermore, the data presented in Figure 4 suggest a saturation of the activity of the expressed proteins with increased fluorescence. This suggests that increased production of the desired target protein does not necessarily lead to a corresponding amount of functional protein. This highlights an indirect problem that is encountered in biotechnological applications for optimal protein expression: apart from the DNA sequence itself, an optimization of the entire expression system and conditions is required.

A possible explanation for this plateauing behavior is that two of codons’ translationally relevant properties could be inversely correlated: more translationally fast codons could be less accurate. Here, accuracy refers to several translational errors: amino acid misincorporation (missense errors) or more severe processivity errors, such as ribosomal drop-off [9, 37]. In such a scenario, the artificial increase of the efficacy of a sequence by means of codon optimization –that is, the enrichment of sequences with fast codons– would, as a side effect, imply the artificial accrual of inaccurate codons in the same sequence. This would ensure high numbers of protein, but the fraction of functional products would be diminished.

One strategy to counteract this problem has previously been introduced by the usage of a dual reporter system, that in addition to protein expression also reports the presence of falsely folded inclusion bodies [38]. This framework could by deployed to address these questions in a more precise manner.

A further concern relates to the sample of the sequences analyzed for the test for epistasis. The set of optimized sequences used was not randomly sampled around a well-known fitness optimum. Therefore, the tests performed for the detection of epistasis may be biased towards above-average expression, if the optimization algorithms were to possess the property that they are capable to raise protein expression with a minimum amount of synonymous changes. However, this does not appear to be a target of the optimization function, since often, the number of synonymous changes proposed by the algorithms range from a minimum of 200 synonymous changes to almost 350 changes that is, given a sequence with a codon length of 360, almost every codon is exchanged in some instances. Furthermore, the test for epistasis does not explicitly state that the sampled sequences around the optimum must be sampled randomly [31, 39].

A last caveat concerns the identity of the optimal sequence for protein expression in the investigation of epistasis. The test for epistasis assumes that the reference strain occupies a local or global optimum, such that all other sequences are separated from the reference sequence by mutations that add a deleterious effect to protein expression. Thus, the misidentification of the best expressing sequence could lead to biases that may affect the test, because if the true best performer sequences are close to the reference strain, a regression curve will have to curve downwards to capture this feature, leading to negative *b* coefficients. To address this issue, we ran several regressions on a subset of possible candidates for best performing sequences, always retrieving our original results. This indicates that the signal for the result stems from the expression characteristics of sequences that are at large Hamming distances from the reference sequences of in the candidate subset, and not the aforementioned possible statistical artefact.

Our results should be interpreted in the context of the broad body of evidence linking codon usage to protein expression and its use for codon optimization. Codon usage has been shown to greatly impact protein expression [11]. However, the variation in obtained gains by codon optimization is large, typically ranging between five to fifteen fold gain in *E. coli* [3, 12].

While several sequence features are accounted for during optimization, such as codon adaptation index, GC-content, avoidance of repeats as well as other motifs (see Table 2 in [3]), the exact nature of many optimization algorithms remains confidential [11, 16]. A large fraction of optimization approaches rely on all-optimal codon strategies [40], but the research field as a whole is moving away from such naïve algorithms [12].

The large variations in protein expression gains as well as in optimization approaches underscores a conspicuous lack of understanding in the mechanisms that give rise to the protein expression level of a sequence [11, 16, 41]. In fact, a recent study achieving a correlation coefficient of 0.76 between measured and predicted expression levels relied on machine learning instead of a mechanistic theory to produce estimates [42]. Reviews report that about 40% of soluble protein targets fail to reach meaningful expression levels [4].

Current limitations to accurately predict protein expression are also reflected by a paucity of predictive theory [4, 42]. The majority of theoretical work on codon usage bias relating to protein expression is aimed at understanding how evolution may shape sequences to maintain a certain protein expression level at minimal energetic expenditure [5, 8, 9, 27, 37, 43, 44], or alternatively, at leveraging statistical associations identified from large data sets to improve predictive quality [45–48].

One of the few instances where protein expression prediction is grounded in a mechanistic foundation, is given in a series of papers by Gilchrist and Shah. These authors have pioneered the approach of explaining codon sequence structure via evolutionary principles [5, 8, 9, 27, 37, 43, 44, 49]. This line of research focuses on determining a function, 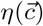, that appropriately represents the cost to benefit ratio of a sequence, 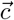. Natural selection works towards minimizing *η*, while being counteracted by mutation (with corresponding, organism-specific biases). The selection pressure exerted upon the sequence towards optimizing its cost-benefit ratio is assumed to be proportional to the target protein expression rate of that sequence. Thus, if the population is assumed to be in mutation-selection balance, the observation of a sequence with a given *η* holds information on its target protein expression rate. Landerer, Gilchrist and Shah have refined Bayesian methods to predict this target protein expression rate given a sequence [37, 43, 49]. However, it is important to note that this method is geared towards predicting target protein expression rates only, that is, the rate or protein expression that is optimal for the organism’s survival in the environment it adapted to.

Our results do corroborate some of the findings of other, more extensive explorations of the synonymous sequence space of a sequence, but are also at odds with some of their findings. For instance, Kudla et al. expressed 154 synonymous sequences of the reporter protein GFP, and found no correlation between codon usage bias and fluorescence levels [35]. Intuitively, this would imply that the strategy underlying most of the codon optimiziation strategies studied here should not have been effective. However, we do observe that the most codon biased sequences –AnaCoda, JCat and GeneGA– do indeed result in the highest expression levels.

Furthermore, the study reported very large increase in fluorescence (i.e., protein expression), of about 250 fold of the original sequence [35]. The two fold increases observed here are much smaller. It is possible that either other variables of the sequence, such as its length and distance from the possible optimum, affect the potential for gain.

Perhaps the most extensive exploration of protein expression patterns in synonymous sequence space has been carried out by Cambray et al. [36]. The authors statistically analyzed the effect of eight factors conjectured to govern protein expression levels of 244’000 sequences in *E. coli*, and concluded that about 50% of variation remains currently unexplained [36]. Codon usage bias did explain around 26% of variation under conditions where translation initiation was facilitated. If such facilitation was absent, codon usage biases contribution was strongly diminished to negligible levels (Fig 2a in [36]). Their analysis suggests that mRNA secondary structures located around the 5’-end exert the strongest influence upon expression levels, accounting for 83% of total effect, in accordance with what was found in later analyses of expression [42, 50, 51].

Corroborating our results, Cambray et al. found evidence for “roughness” of the sequence expression landscape [36], an expected consequence of epistatic effects. The authors concluded that this suggests the presence of unknown sequence descriptors containing important predictive information. We hypothesize that such roughness may arise from epistasis: Individual codon contributions may add up disproportionately, depending on codon position and genetic background [29]. A possible mechanism for such epistasis was identified by Cannarozzi et al. [52], which found that clusters of similar codons appear to enhance translation efficiency by facilitating the reusage of the same tRNA. Thus, the presence of the same codon upstream may enhance the contribution to expression of a codon beyond what would be expected if that first codon was missing, giving rise to epistasis.

The mirroring techniques developed at ICLS are similar in spirit and implementation to the *codon harmonization* techniques pioneered by Angov et al. [53]. Similarly to the mirroring technique adopted here, the rationale behind codon harmonization is the conjecture that clusters of non-optimal codons serve the purpose of aiding the nascent protein to fold correctly [54]. To optimize the source sequence, they first established a set of codon usage frequencies for each amino acid residue in the source organism’s genome. An analogous list was then produced for the host organism. Original codons where replaced by hast codons that most closely matched usage frequency [53]. Link/end segments are treated by a special procedure. Unlike the mirroring technique used here, codon harmonization thus relies on usage frequencies instead of translational inefficiency estimates (inferred by the AnaCoda package) to inform the codon replacements. Using this technique, Angov et alṡuccessfully codon harmonized the sequence for MSP1 (FVO) from *Plasmodium falciparum* -a candidate antigen for a Malaria vaccine- and achieved a sixty-fold increase in protein yield when expressed in *E. coli*. Our results differ starkly the mirrored sequences were among the least expressed. However, the poor expression attained by the mirrored and capped sequence, where all of the most inefficient codons were replaced by the second most inefficient codons, suggests that the rationale underpinning this approach warrants more detailed investigation [55].

The quest for understanding protein production is ongoing [16] and the question whether there exist fundamental limitations to its predictability remains to be answered conclusively. But a more profound understanding of the underlying mechanisms of protein synthesis holds the promise to acquiring the capacity to engineer synthetic organisms, for instance via designed genetic codes [56].

## Materials and Methods

### Data

#### Plasmid collection and strain preparation

All DNA sequences for expression were designed as fusion proteins, containing a C-terminal linker with the GFP11 fragment that allows detection of expression in a split-GFP assay. Cloning and ligation of the wild type sequence of AtCAD5 into pET28b was performed using standard restriction and ligation techniques. The wild type sequence of AtCAD5 was supplied as gene block from Integrated DNA Technologies (IDT). NdeI and XhoI sites were used for the restriction, followed by ligation by the T4 DNA ligase of the gel-purified DNA fragments. Correct insertion of the wild type sequence of AtCAD5 was confirmed by sequencing. Remaining DNA sequences for AtCAD5 were ordered at Twist Bioscience as clonal genes in pET28b, whereas a selection of constructs was generated by adapting specific codons to the desired codon sequences (Supplementary Table 1). To that end, a PCR using a primer pair that contains the desired nucleotide change was performed, followed by digestion of the template strand by DpnI (see *Supplementary Information* on codon optimized sequences). Correct plasmid sequences were confirmed by sequencing. Plasmids were transformed into chemically competent *E. coli* BL21(DE3) or electrocompetent *E. coli* K12(DE3) cells for expression. *E. coli* K12(DE3) corresponds to the K12 strain MG1655 ΔendA ΔrecA (DE3) and was a gift from Kristala Prather (Addgene plasmid # 37854)[57].

#### Preparation of Split-GFP assay detector fragment GFP1-10

The previously reported DNA sequence for the GFP1-10 detector fragment was obtained as gene block from IDT [58]. Amplification and elongation with flanking NdeI and HindIII restriction sites was performed by PCR. The amplified fragment and pET22b were digested, DNA fragments were gel purified and ligation using the T4 DNA ligase was performed. Correct insertion of the GFP1-10 fragment into pET22b was confirmed by sequencing. Expression was performed in *E. coli* BL21(DE3), with all media being supplemented with 100 *μ*g/ml ampicillin. A 10 L fermentation in a 15 L reactor was conducted in TB (terrific broth) medium. Inoculation was performed 1:20 using a TB overnight culture grown at 37 ^°^C, 180 rpm. Cultivation parameters were kept at 37 ^°^C, a pH range of 7.0 – 8.5 and *p*_O2_ *>* 35%. Expression of GFP1-10 was induced with 100 *μ*M IPTG at OD_600_ 5.5, continuing cultivation for 22 h. The biomass was pelleted by centrifugation at at 3428 x g, 4 ^°^C for 20 min and stored at −20 ^°^C until further usage. The purification of GFP1-10 from inclusion bodies was achieved based on previously established protocols [28, 58]. Cell pellets were resuspended in 15 ml per 5 g wet biomass of TNG buffer (100mM Tris-HCl pH 7.4, 100 mM NaCl, and 10% glycerol) and incubated on ice for 5 min. Cell disruption was performed in three cycles by ultrasonication (2 min, amplitude 50%, pulse 1s/1s). The inclusion bodies were pelleted by 30 min centrifugation at 19000 x g and 4 ^°^C, followed by 6 cycles of resuspension in 10 ml TNG buffer and 10 min centrifugation at 3428 x g and 4 ^°^C. The final centrifugation step was performed at 16000 x g at room temperature, and the pellet was stored at −80 °C until further usage. The inclusion bodies were resuspended in 1 M urea to a concentration of 75 mg/ml. Following 1 h agitation at 600 rpm at room temperature, the solution was centrifuged for 1 min at 16000 x g, before transferring the supernatant to 25 ml TNG buffer. Concentration of GFP1-10 was determined by a NanoDrop One UV-Vis Spectrophotometer (280 nm, 24.443 kDa, 17.545 l mol^−1^ cm^−1^) and adjusted to 0.4 mg/ml with TNG buffer. Aliquots were stored at −20 ^°^C until further usage.

#### Protein expression for lysate preparation

Growth of *E. coli* BL21(DE3) and *E. coli* K12(DE3) for expression of codon variants was performed in quadruplicates in 96-well deep well plates in two independent experiments, resulting in a total of eight replicates per strain. All media were supplemented with 50 *μ*g/ml kanamycin. Overnight precultures in 900 *μ*l LB medium were grown at 37 °C, 300 rpm. Precultures were used to inoculate 1200 *μ*l fresh TB medium in a 1:1000 ratio. Induction with 100 *μ*M IPTG was performed in early-mid exponential phase at OD_600_ 1.21 ± 0.27 or 0.75 ± 0.08 for BL21(DE3) or K12(DE3) respectively, followed by expression at 20 ^°^C. After 20 ± 2 h, the well volume was adjusted to 900 *μ*l and the plate was centrifuged for 20 min at 3428 x g at 4 ^°^C. The supernatant was discarded, and cell pellets were stored at −20 ^°^C until further usage. For preparation of lysates, each pellet was resuspended in 180 *μ*l of BugBuster^®^ Master Mix supplemented with 3.23% (v/v) Protease Inhibitor Cocktail Set II. For strains with lower end OD - mostly affecting K12(DE3) strains that generally grow lower than BL21(DE3) - the resuspension volume was reduced to maintain comparable biomass concentrations of 0.083 ± 0.030 OD/*μ*l in the lysate. Following incubation for 30 min at room temperature, the plate was centrifuged for 20 min at 3428 x g at 4 ^°^C. The supernatant was transferred to a fresh plate for analysis of protein expression and activity.

#### Split-GFP assay

In black 96-well microtiter plates with clear bottom, 12.5 *μ*l cell lysate from deep well plate expression were mixed with 187.5 *μ*l GFP1-10 detector fragment. The plate was sealed and incubated at 32 ^°^C and 100 rpm for 20 h, to allow stable formation of fluorescent GFP-complexes signal. Fluorescence was measured with a Tecan Infinite^®^ 200 PRO plate reader, using extinction at 485 nm and emission at 520 nm. Within each assay plate, the average of the background GFP signal using the empty pET28b vector was subtracted from all other strain signals. The average and the standard deviation of all replicates was determined. Replicates that deviated more than 1.5 x standard deviations from the mean were considered outliers and were subsequently excluded from calculations.

#### Enzymatic detection of AtCAD5 activity

Enzyme activity was determined using clear 96-well microtiter plates with clear bottom similarly as previously reported in [59]. The reaction mixture for the assessment of enzyme activity in the lysate consisted of a final concentration of 485 *μ*M NADPH and 2.4 mM cinnamal aldehyde in 0.1 M Sodium Potassium Phosphate buffer (pH 6.25). A fresh 30x dilution of cell lysate from deep well plate expression was prepared in 0.1 M Sodium Potassium Phosphate buffer (pH 6.25). To initiate the reaction, 10.0 *μ*l of the diluted lysate was added to the reaction mixture. NADPH consumption was followed by A_340nm_ measured with a Tecan Infinite^®^ 200 PRO plate reader over at least 2 min in 11.7 sec intervals. To assess AtCAD5 activity in the lysate, the inital slope of the reaction was determined. The average and the standard deviation of all replicates was determined. Replicates that deviate more than 1.5 x standard deviations from the mean were considered outliers and were subsequently excluded from calculations.

#### Genomic data for A. thaliana and E. coli used for Codon Optimization

In this study, we used two genome data sets – one for *Arabidopsis thaliana* (*A. thaliana*) and one for *E. coli* – as well as a protein expression data set – for *E. coli* – to calibrate our codon optimization model via the AnaCoda package.

For *A. thaliana*, we downloaded the coding sequences from the Arabidopsis thaliana information resource (TAIR) [60]. These sequences contained in the Araport11 blastsets, file Araport11_cds_20210622.fasta. The wildtype sequence for the gene AtCAD5 was also downloaded from TAIR, reference At4g34230.

For *E. coli*, we obtained the coding sequence data for the strain K12, substrain MG1655, from the National Center for Biotechnology Information (NCBI), accession number NC 000913.3. The data include 4319 protein coding sequences (see https://www.ncbi.nlm.nih.gov/data-hub/taxonomy/511145/ [61–64]).

#### Protein expression level data for E. coli

Protein expression data for *E. coli* were generated in [7], and were converted into a useful format in the publication [49], and can be found on the Github project https://github.com/acope3/Signal_Peptide_Scripts/blob/master/Data/Empirical/ROC_Phi/Ecoli_K12_MG1655_main_phi.csv. Briefly, to convert the data from [7], the authors of [44] utilized the mRNA translational efficiency values in the Supplementary Table S4 in [7], and multiplied them with the corresponding mRNA levels of the same gene to obtain the protein expression level for that gene. These levels were then normalized in such manner that the average expression level is one.

#### Codon-specific elongation rate data for E. coli

In this study, we used three different sets of codon-specific elongation rate data for*E. coli*. The first set was estimated by Shah and Gilchrist [27] mostly on the basis of tRNA abundancies. They estimated cognate elongation rates of codons by combining gene copy numbers of cognate tRNA (as proxies for cognate tRNA abundances) and probabilities of elongation of near- and pseudo cognates. The table with the codon-specific elongation rate values (column *R*_*c*_) was retrieved from Table S2 of the Supporting Information of [27].

The second set of codon-specific elongation rates was derived from elongation time estimates. Proxies of elongation times were estimated via de AnaCoda package’s ribosomal overhead cost (ROC) model, which assumes that the selection coefficient of a synonymous sequence variant is directly proportional to the overall elongation time of the sequence [22, 43]. To obtain a table of relative codon-specific elongation times, we ran AnaCoda’s ROC model on the full *E. coli* genome with normalized protein expression level data. The values in the table of estimates corresponds to codon-specific overhead costs, which are assumed to be proportional to the elongation time estimates for each codon. All the overhead costs within a synonymous codon family are given relative to the least costly codon within the family, which by default has a cost of zero.

A third data set by Rudorf and Lipowsky was estimated on the basis of ribosome occupancy data [26]. The authors employed a Markov model to infer codon-specific elongation rates under different sets specific bacterial growth rates, assuming a 2-1-2 pathway of tRNA release at the E site. Their estimates are found in the S2 Table of the Supporting Information of [26]. We used the estimates for a specific growth rate of 0.7 per hour, which is close (within 0.2 per hour) to the growth rate measured for E. coli in the experimental setup of this study for all replicates examined.

#### Summary of external data sources

Table II gives a summary of the data used from external sources, including coding sequences for the gene to be optimized, genomic data, and especially data on protein expression levels in *E. coli* and codon-specific translation information for *E. coli* as well.

### Codon Optimization of the Sequences

#### Anacoda-based all-optimal codons strategy for sequence design

For the design of the codon optimized sequence termed *AnaCoda*, we first ran the ribosome overhead cost (ROC) model [27, 43] of the AnaCoda package [22] on the *E. coli* genome, including information on normalized protein expression levels measured during *E. coli* ‘s exponential growth phase. This regression yielded estimates for both, codon-specific mutational biases as well as codon-specific biases in the cost of translation as would by inferred if these were the main targets of selection for synonymous codon change.

Under the ROC model, the cost of translation corresponds exactly to the sum of the elongation times of all codons of the sequence times a factor to convert these into selection coefficients. Thus, the codon-specific cost estimates from AnaCoda are directly proportional those codons’ elongation times [27, 43]. As a caveat, the translational costs can only be computed relative to the least costly (that is, most time-effective or translationally fast) codon within a codon family for the same amino acid.

With these preparations, we proceeded to replace each codon of the AtCAD5 wildtype sequence with the optimal codon of its family as estimated under the ROC model. That is, each codon at every position was replaced by a codon that under the ROC model has a translational cost of zero, and thus the smallest elongation time within its codon family. Notably, the optimized sequence differs from other sequences optimized by other, more classical codon optimization algorithms. This is because for the amino acid threonine (T), AnaCoda identifies the codon ACT as optimal, while JCat identified the codon ACC as optimal.

#### Proprietary sequences of CAD5

To obtain codon optimized sequences from the commercial providers, we utilized the optimization algorithms freely available on the providers’ official websites. Table III gives a summary of the urls where these proprietary algorithms can be accessed. If the optimization algorithms worked by stochastic optimization principles, we repeated the optimization three times. All optimization algorithms allowed for the exclusion of the restriction sites BamHI [GGATCC], EcoRI [GAATTC], EcoRV [GATATC], NdeI [CATATG] and XhoI [CTCGAG], necessary for plasmid construction.

#### Anacoda-derived mirroring optimization strategies

To mirror the original elongation rate pattern in the new organism, we first had to assess codons’ translational efficiencies in their original organism’s genetic environment. We ran a regression of the ROC model of the AnaCoda package [22, 23] on the *A. thaliana* genome, yielding tables of relative codon-specific elongation times for the *A. thaliana* genome.

Subsequently, the same regression procedure was performed for *E. coli* (additionally taking protein expression levels as input), yielding an analogous, codon-specific translational efficiency table.

The wildtype sequence was then optimized as follows: Each original codon was replaced by a codon that occupied the same relative efficiency rank within its synonymous codon family, thus replacing the most inefficient original codons with the most inefficient new codons, and so forth. In the publication, we term this the *mirrored* sequence. The code detailing this algorithm is available in the *Supporting Information*.

In a second approach, the mirrored sequence was pruned of restriction sites (see section *Proprietary sequences of CAD5* for a list of the restriction sites). When removing restriction sites, we first identified the codons with the most similar translational efficiency of a synonymous codon family member for each of the two codons on the site. We then replaced that codon with its most similar synonymous counterpart.

In a third approach, we replaced all maximally inefficient codons with the second most inefficient codons of their respective synonymous codon family, thereby “capping” inefficiencies in the sequence. We term this approach *mirrored and capped*.

#### Summary of the codon-optimized sequences

Table III gives a summary of the optimized sequences that were heterologously expressed in this study. Table also gives reference to the codon optmization algorithms employed by commercial providers of the gene products, as well as the references, if found, of the scientific publications describing the optimization algorithms.

### Multidimensional Scaling

Efficiency profiles may be interpreted as vectors in which the *i*^th^ entry corresponds to the elongation time of the codon at the *i*^th^ position of that sequence. This allows to define a similarity metric between sequences and subsequently to apply MDS, a technique that allows for the representation and visualization of multidimensional similarity matrices in a two-dimensional Cartesian space, while preserving inter-element relationships –as expressed via a cost function termed *stress* to be minimized– as much as possible [70, 71] (see Materials and Methods).

The distance metric used in Figure 2 is the Euclidian distance 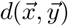 between relative elongation time or elongation rate profiles, given by

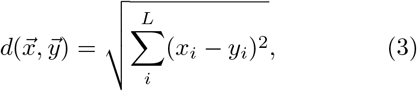

where 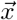 and 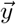 are either the elongation time or elongation rate vectors of two sequences, and where *x*_*i*_ and *y*_*i*_ are the elongation time or rate of a codon at the i^th^ position of the sequence, respectively. Additionally, *L* is the length of the sequence in codons.

For Figure 2A), the elongation time profiles were derived by using relative translation inefficiencies inferred under the ribosomal overhead cost (ROC) model of the AnaCoda package [22, 23, 43].

The distance metric used in Figure 2B) was the Euclidian distance between the elongation rate profiles of the sequences derived from the Shah data set of codon-specific elongation rates [27], while for C) they were derived from the Rudorf data set [26].

For all MDS procedures, we used Kruskal’s non-metric multidimensional scaling [70, 71].

## Supporting information

Protein activity data

All raw data

sequence names mRNA

## Acknowledgments

Victor Garcia was supported by SystemsX.ch – the Swiss Initiative for Systems Biology [grant number 51FSP0_163566], the Swiss National Science Foundation [grant number 31003A_182330] and the Digitalization Initiative of the Zurich Higher Education Institutions (DIZH fellowship grant). Marco Gees was supported by the Swiss National Science Foundation [grant number 31003A_182330]. Marco Gees, Zrinka Raguz Nakic, Victor Garcia and Christin Peters thank the Zurich University of Applied Sciences for the internal funding of the project “Die Exploration der Gen-Harmonisierung”.

